# Multi-locus pairing expands the functional repertoire of two-peptide bacteriocins

**DOI:** 10.64898/2026.11.12.766112

**Authors:** João Lança, João Borralho, Raquel Sá-Leão

**Affiliations:** Laboratory of Molecular Microbiology of Human Pathogens, Instituto de Tecnologia Química e Biológica António Xavier, Universidade Nova de Lisboa, Oeiras, Portugal

**Author notes:** Corresponding author: Raquel Sá-Leão. Co-first authors (João Lança, João Borralho).

## Abstract

Two-peptide bacteriocins are generally viewed as self-contained systems whose peptides are encoded at the same biosynthetic locus and co-expressed to form an active antimicrobial pair. Here, we show that functional pairing extends beyond this organization. Comparative genomic analysis revealed frequent co-occurrence of putative two-peptide bacteriocins encoded at distinct chromosomal loci in streptococci. Testing pairwise combinations of 47 peptides produced by cell-free protein synthesis identified 54 combinations with enhanced antimicrobial activity, with inter-locus combinations more frequent than intra-locus combinations. In *Streptococcus mitis* strain C22, four dispersed bacteriocin loci are coordinately expressed during exponential growth despite regulation by partially independent signaling pathways, providing an opportunity for inter-locus pairing. Split-luciferase complementation assays showed that active peptide pairs associate and that these interactions depend predominantly on amino acid sequence. Using a bacteriocin-free *S. mitis* background, we further show that components produced by distinct bacterial cells can act together to increase producer competitiveness in co-culture with *S. pneumoniae*. These findings reveal a modular organization in which peptides encoded at different loci can form functional combinations, including when produced by separate cells. This flexibility expands the antimicrobial repertoire available to individual strains and provides a mechanism for bacteriocin diversification while retaining functional compatibility.

## Introduction

Bacteriocins are ribosomally synthesized antimicrobial peptides that mediate competition between bacteria. By selectively inhibiting competitors, bacteriocins can influence community composition^1^, niche occupation^2–4^, and horizontal gene transfer^5,6^. Advances in genome sequencing and biosynthetic gene cluster mining have revealed that bacteriocins constitute a widespread and remarkably diverse class of antimicrobial molecules^7–9^.

Streptococci are particularly rich in bacteriocin systems, and genome mining has identified thousands of putative bacteriocin peptides, including lantibiotics, circular, and linear non-modified bacteriocins^6,9–13^. In *Streptococcus pneumoniae*, the bacteriocin-like peptide (*blp*) locus is among the most polymorphic regions of the chromosome and can encode a diverse repertoire of bacteriocins and immunity proteins^14^. Likewise, comparative genomic analyses of *Streptococcus mitis* have identified more than 90 unique bacteriocin peptides within competence-associated bacteriocin (*cab*) loci^6^. Together, these observations indicate that diversification of bacteriocin repertoires is a pervasive feature of streptococcal evolution.

Despite the large number of bacteriocins predicted by genome mining, the functional significance of this diversity remains poorly understood. A major obstacle is that only a small fraction of predicted bacteriocins has been experimentally validated^7,15–17^. Functional characterization is often hindered by difficulties in heterologous expression arising from peptide toxicity and low yields, as well as by frequent co-production of multiple bacteriocins by native producer strains. In addition, limited knowledge of the environmental and regulatory conditions governing bacteriocin expression, together with the difficulty of identifying their natural target organisms, has impeded efforts to define the biological functions and activity spectra of individual bacteriocins.

These challenges are particularly critical for two-peptide (class IIb) bacteriocins, whose antimicrobial activity depends on the cooperative action of two distinct peptides^18,19^. These systems are regarded as genetically self-contained, with compatible peptide partners encoded within the same biosynthetic locus and co-expressed as cognate functional pairs^18^. Although isolated studies have shown that a small number of two-peptide bacteriocins can tolerate some non-cognate peptide combinations in vitro^20–22^, whether such observations represent exceptional cases or a broader organizational principle remains unknown. This uncertainty raises a fundamental evolutionary question: how is functional compatibility maintained as bacteriocin repertoires diversify? If productive pairing requires strict peptide specificity, diversification should be constrained by the need to preserve cognate peptide interactions. Alternatively, bacteriocin peptides may exhibit greater flexibility in pairing than previously appreciated, enabling the emergence of new functional combinations while facilitating the expansion of bacteriocin repertoires.

Here, we show that positive co-occurrence of bacteriocin peptides encoded at distinct chromosomal locations is frequent across streptococcal genomes. Using cell-free protein synthesis, we systematically mapped the functional interaction landscape of 47 bacteriocin peptides from seven streptococcal strains and identified extensive functional pairing, including numerous combinations between peptides encoded at different chromosomal loci. We further show that bacteriocin loci distributed across the chromosome can be coordinately expressed in *S. mitis*, providing an opportunity for compatible peptides to be produced concurrently despite their distinct regulatory contexts. Split-luciferase complementation assays demonstrated that active peptide pairs associate, with interactions predominantly dependent on primary peptide sequence. Finally, using a bacteriocin-free *S. mitis* strain, we show that individual bacteriocin components produced by distinct bacterial strains can cooperate to increase competitiveness against *S. pneumoniae*. Together, these findings establish a modular view of two-peptide bacteriocins in which functional pairing is not restricted to genetically linked partners or peptides produced within the same strain. Instead, bacteriocin peptides can participate in broader compatibility networks that expand the functional repertoires of their hosts.

## Results

### Comparative genomics reveals widespread co-occurrence of bacteriocins encoded at distinct chromosomal loci

We hypothesized that functional two-peptide bacteriocins could arise from peptides encoded at distinct chromosomal loci rather than exclusively from genetically linked genes. To test this hypothesis, we examined whether streptococcal genomes frequently encode multiple dispersed bacteriocins, and whether bacteriocin genes from different loci are preferentially maintained together.

We analysed a recently generated catalogue of streptococcal bacteriocins identified by large-scale genome mining of RefSeq genomes^9^. This dataset comprises more than 9,000 streptococcal genomes encoding over 3,000 unique bacteriocins (**Fig. 1a and Supplementary Table 1**). To focus on putative two-peptide bacteriocins capable of inter-locus pairing, we excluded peptides located near contig boundaries, retained only bacteriocins containing (G/A/S)xxx(G/A/S) motifs (frequent in two-peptide bacteriocins^23–25^), and defined distinct loci as bacteriocins separated by at least 30 kb. We then quantified bacteriocin repertoires within individual genomes and tested for significant co-occurrence between peptides encoded at different loci (**Supplementary Table 2**).

**Figure 1.**
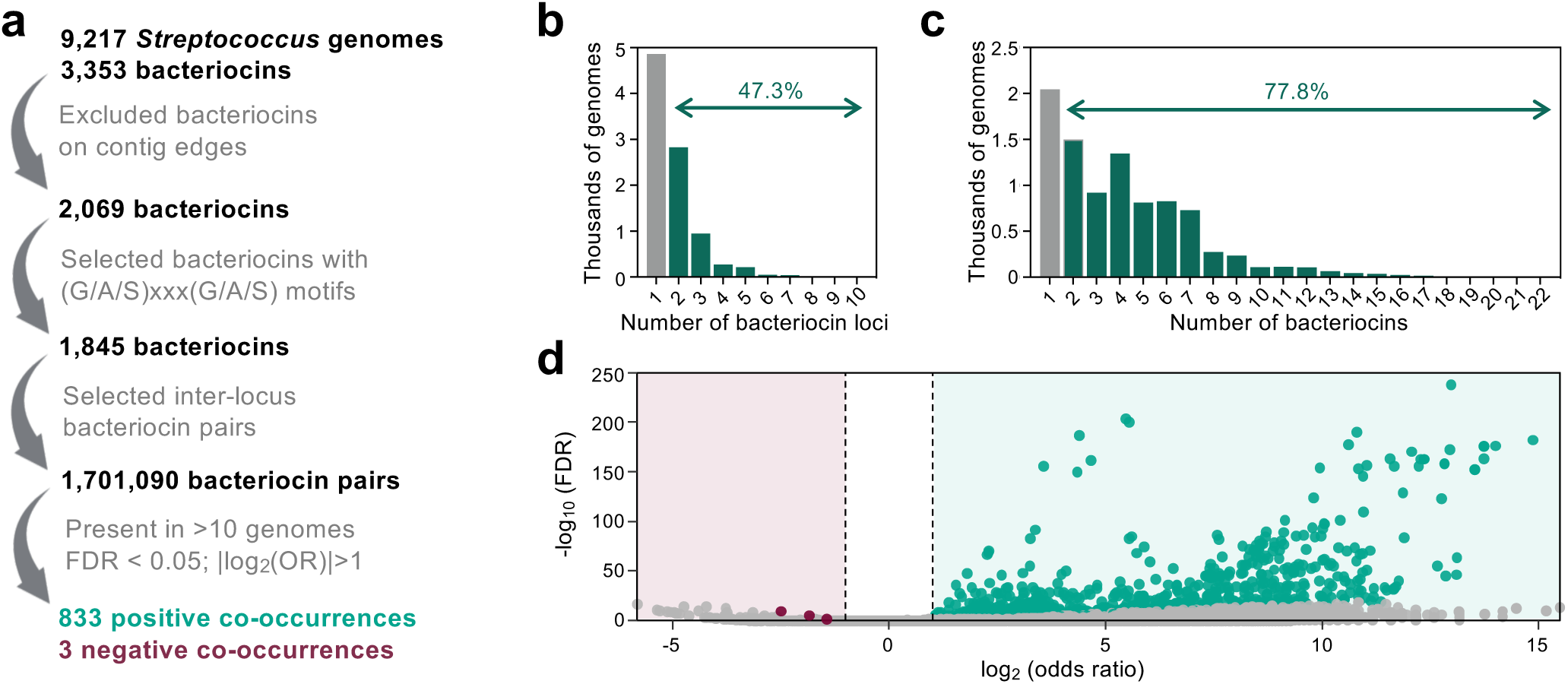
**Comparative genomics reveals co-occurrence of bacteriocins encoded at distinct chromosomal loci**. (a) Workflow of the comparative genomic analysis. Bacteriocins identified in a previous large-scale genome mining study^9^ were first filtered to exclude peptides located within 30 kb of contig edges. Bacteriocins containing (G/A/S)xxx(G/A/S) motifs in their predicted mature sequences were then retained, and bacteriocin loci were defined using a minimum distance of 30 kb between bacteriocins. Finally, pairwise co-occurrence analysis was performed for bacteriocins encoded at distinct chromosomal loci within individual genomes. Associations were considered significant when false discovery rate (FDR) < 0.05, an absolute odds ratio (OR) |log_2_(OR)| > 1, and a bacteriocin pair was found in at least 10 genomes. (b,c) Distribution of the number of (b) bacteriocin loci and (c) bacteriocins containing (G/A/S)xxx(G/A/S) motifs per streptococcal genome. (d) Pairwise co-occurrence analysis of bacteriocins encoded at distinct chromosomal loci across streptococcal genomes. Significant positive and negative associations are shown in green and pink, respectively. Grey datapoints with FDR < 0.05 and |log_2_(OR)| > 1 represent bacteriocin pairs detected in fewer than 10 genomes and therefore excluded. Statistical analysis was done using Fisher’s exact test and *P* values were adjusted for multiple comparisons using the Benjamini-Hochberg FDR.

Distributed bacteriocin repertoires were common across streptococci. Nearly half of genomes (47.3%) encoded bacteriocins at two or more locations (**Fig. 1b**), and 77.8% encoded at least two distinct (G/A/S)xxx(G/A/S)-containing bacteriocins (**Fig. 1c and Supplementary Fig. 1**), indicating that many genomes possess the potential for non-cognate bacteriocin pairing. We next asked whether peptides encoded at distinct loci are preferentially maintained together during evolution. Among bacteriocin pairs that met our criteria, significant positive associations greatly outnumbered negative associations (**Fig. 1d**), indicating that several bacteriocin pairs encoded at different locations frequently co-occur across streptococcal genomes.

Together, these comparative genomic analyses reveal ample opportunities for inter-locus bacteriocin pairing and motivated us to experimentally determine whether peptides encoded at distinct locations can assemble into functional antimicrobial pairs.

### Systematically pairwise in vitro screening reveals extensive inter-locus pairing among streptococcal bacteriocins

We recently identified seven streptococcal isolates with potent antagonistic activity against *S. pneumoniae* (*S. oralis* A22 and *S. mitis* strains B22-G22)^13^. Most bacteriocins encoded by these strains are absent from pneumococcal genomes, and we have observed that *S. pneumoniae* is typically susceptible to their antimicrobial activity^6,13,16,17^. In addition, their complete genome sequences are available enabling unambiguous assigment of bacteriocins to individual chromosomal loci^6,13,16,17^. Together, these features make this collection a suitable system for systematically investigating functional pairing of bacteriocins.

To determine whether bacteriocins encoded within the same genome can form functional pairs, we produced 47 putative bacteriocin peptides containing (G/A/S)xxx(G/A/S) motifs^23–25^ using cell-free protein synthesis. This corresponded to all the bacteriocins collectively encoded by the seven strains excluding those predicted to have post-translational modifications (which we were unable to produce by cell-free protein synthesis) and those with high prevalence in pneumococci (e.g. CibAB)^13^. When tested individually against *S. pneumoniae* D39, only six peptides (12.8%) exhibited detectable antimicrobial activity, indicating that most peptides lacked measurable activity under these conditions (**Fig. 2a**).

**Figure 2.**
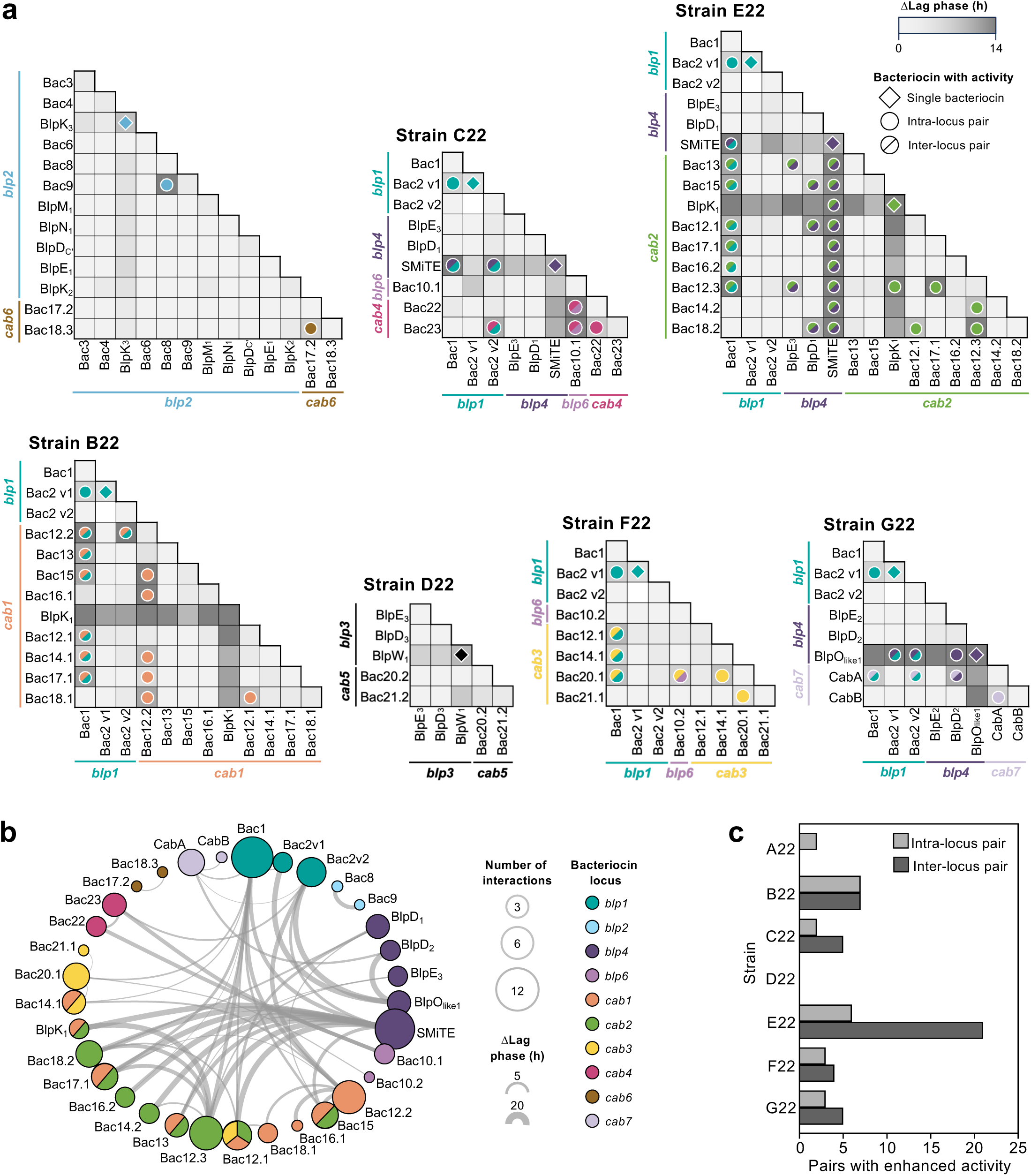
Systematic functional screening identified extensive inter-locus bacteriocin pairing. (a) Pairwise antimicrobial activity of 47 putative bacteriocins containing (G/A/S)xxx(G/A/S) motifs encoded by seven streptococcal genomes (of strains S. oralis A22, and S. mitis B22, C22, D22, E22, F22, and G22). Mature bacteriocins were produced by cell-free protein synthesis and tested individually and in pairwise combinations against S. pneumoniae D39. Bacteriocins are grouped according to chromosomal locus (previously annotated13). Individual bacteriocins were considered active when delaying exponential growth by ≥ 2 h. Bacteriocin pairs were considered active when delaying exponential growth by ≥ 2 h compared to the effect of either of the bacteriocins alone. (b) Network of active bacteriocin pairs identified in the screening. (c) Distribution of synergystic bacteriocin pairs according to whether both peptides originate from the same chromosomal locus (intra-locus) or from different loci (inter-locus).

We next tested every pairwise combination of peptides encoded within each genome (295 combinations in total). Strikingly, 54 combinations (18.3%) exhibited substantially greater antimicrobial activity than either constituent peptide alone (**Fig. 2a, Supplementary Table 3, and Supplementary Fig. 2**). This included the previously characterized two-peptide bacteriocin CabA-CabB^6^, validating the screening strategy.

The resulting interaction network revealed marked differences in partner specificity (**Figs. 2a, b, Supplementary Table 3**). Some peptides, including Bac8 and Bac9 from the *blp2* locus, formed active pairs with only a single partner. In contrast, Bac1 from the *blp1* locus and SMiTE from the *blp4* locus displayed extensive promiscuity, generating functional combinations with multiple peptides encoded across different chromosomal loci and streptococcal strains.

Surprisingly, productive pairs were identified more frequently between peptides encoded at different chromosomal loci (64.8%) than between peptides originating from the same locus (35.2%) (**Fig. 2c, Supplementary Table 3**). These findings indicate that productive bacteriocin pairing is not constrained by genomic organization and suggest that compatible peptide partners can emerge from distributed bacteriocins repertoires rather than exclusively from genetically linked bacteriocins.

Together, these results reveal an extensive network of functional interactions among streptococcal bacteriocins and demonstrate that inter-locus pairing substantially expands the repertoire of active two-peptide bacteriocins encoded within individual genomes.

### Coordinated expression enables inter-locus bacteriocin pairing in *Streptococcus mitis*

The discovery of numerous functional inter-locus bacteriocin pairs in vitro raised the question of whether peptides encoded at distinct locations are produced simultaneously in their native hosts, thereby creating the opportunity for inter-locus pairing and cooperative activity.

To address this question, we focused on *S. mitis* C22, which encodes four bacteriocin loci (*blp1*, *blp4*, *blp6*, and *cab4*) that collectively give rise to multiple functional inter-locus pairs (**Figs. 2a-c and 3a, and Supplementary Table 3**). While *blp* loci are regulated by BlpC pheromone signaling^13,26–29^, the *cab* locus is induced by competence-stimulating peptide (CSP) through the competence regulatory pathway^6,30^. In C22, *blp1* and *blp4* encode their own cognate pheromones (BlpC1 and BlpC_4_, respectively), whereas *blp6* lacks an associated pheromone gene. *comC,* encoding for CSP_C22_, is located approximatelly 60 kb upstream of *cab4* (**Fig. 3a**).

**Figure 3.**
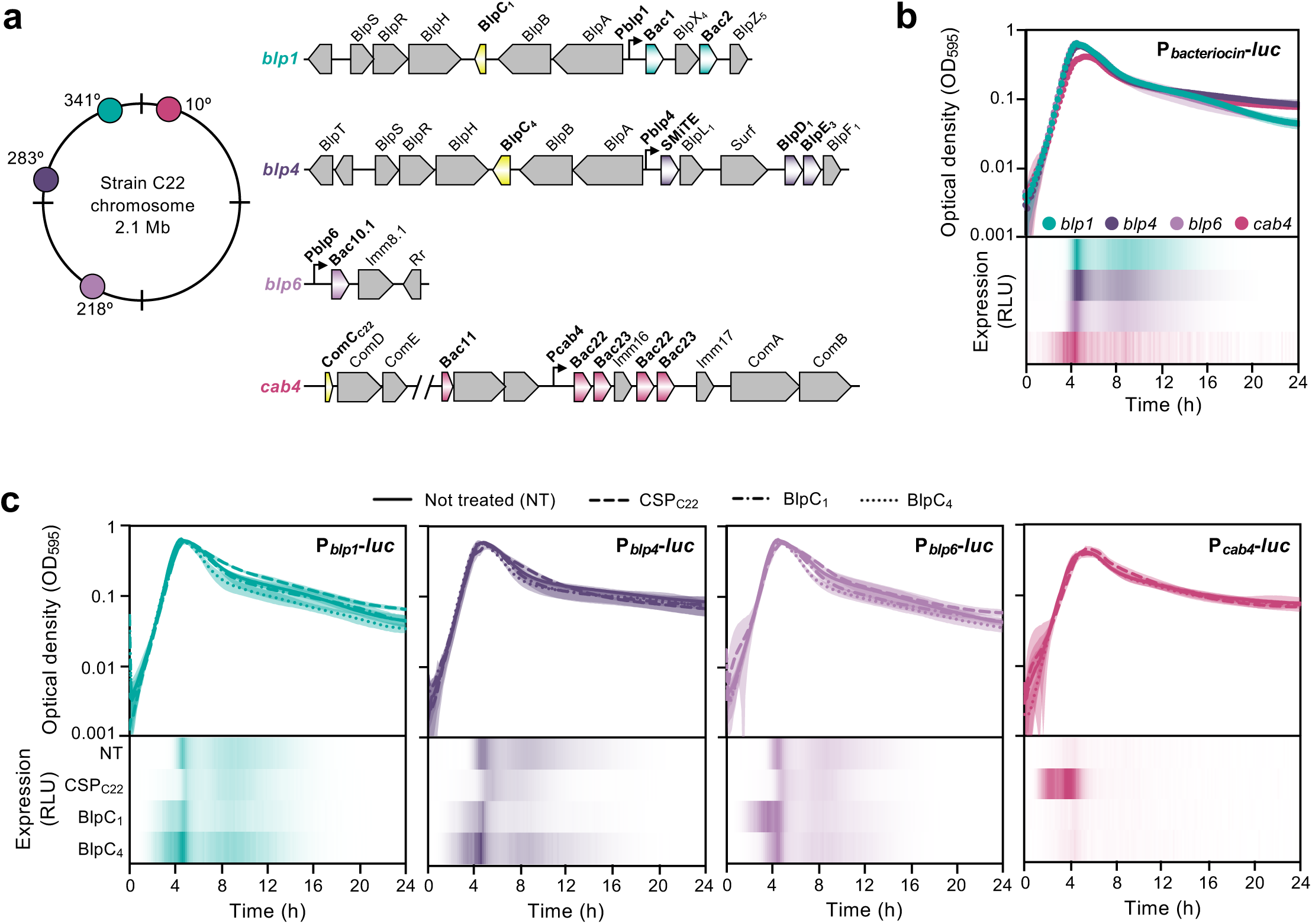
Expression of bacteriocins from distinct chromosomal loci in *Streptococcus mitis*. (a) Genomic organization and location of the four bacteriocin loci (*blp1*, *blp4*, *blp6* and *cab4*) in *S. mitis* C22. Inducing pheromones are shown in yellow. The *comCDE* operon, from which CSP_C22_ is expressed, if found ∼60 kb upstream of *cab4*. Promoters used for luminescence transcriptional reporters are shown (P*_blp1_*, P*_blp4_*, P*_blp6_* and P*_cab4_*). (b) Temporal expression of the four bacteriocin loci during growth. Promoter activity was monitored using firefly luciferase transcriptional reporters fused to the promoter region of each bacteriocin locus. Data represents the mean ± s.d. from three biological replicates, each with three technical replicates. (c) Induction of bacteriocin loci by exogenously added synthetic pheromones. Reporter strains were exposed to 1 μM of BlpC_1_ BlpC_4_, CSP_C22_ or water (not treated, NT) and luminescence was measured throughout growth. Data represents the mean ± s.d. from three biological replicates, each with three technical replicates.

To determine whether these genetically dispersed loci are expressed simultaneously, we generated transcriptional reporter strains (one per locus) in which the promoter from each bacteriocin immunity region drived the expression of firefly luciferase (*luc*). Despite being controlled by distinct signalling pathways, all four loci displayed highly similar temporal expression profiles, with expression peaking during exponential growth (**Fig. 3b and Supplementary Table 4**). Thus, bacteriocins encoded at distinct chromosomal loci are coordinately expressed.

We next investigated whether regulatory cross-talk between pheromone signalling pathways could contribute to this coordinated expression. Both *blp* and competence signalling are regulated by extracellular pH, with acidic conditions inhibiting induction^6,31^. However, we were unable to identify an extracellular pH at which all four bacteriocin loci were transcriptionally inactive in *S. mitis* C22 (**Supplementary Fig. 3 and Supplementary Table 5**). We therefore performed subsequent induction experiments at pH 7.4, a condition in which all four loci were expressed while bacterial growth remained unaffected. Addition of synthetic BlpC pheromones revealed extensive cross-regulation among *blp* loci (**Fig. 3c and Supplementary Table 4**). Both BlpC_1_ and BlpC_4_ activated expression from the *blp1* and *blp4* promoters, while *blp6* was strongly induced by BlpC_1_ while lacking a cognate pheromone gene. In contrast, *cab4* expression was induced exclusively by CSP_C22_ and remained unresponsive to either BlpC pheromone, indicating that competence and *blp* signalling remain distinct under these conditions.

Together, these findings show that bacteriocin loci distributed across the chromosome are coordinately expressed despite being controlled by partially independent regulatory pathways. Cross-regulation among *blp* loci likely contributes to this coordinated expression, whereas *cab4* remains specifically linked to competence signalling under the conditions tested. The temporal overlap in expression provides an opportunity for peptides encoded at distinct chromosomal locations to be produced concurrently, creating the potential for functional inter-locus bacteriocin pairing.

### Synergistic bacteriocin pairs exhibit selective peptide association

The frequent enhanced activity observed among bacteriocin pairs raised the question of whether enhanced antimicrobial activity arises from association between compatible peptide partners or independent additive effects.

We first asked whether peptide compatibility could be predicted computationally by structural modeling of all 295 bacteriocin combinations with AlphaFold 3. Although active pair displayed a slightly higher interface predicted template modeling (ipTM) scores, this difference was not statistically significant (**Supplementary Fig. 4 and Supplementary Table 3**), indicating that AlphaFold could not reliably distinguish active from inactive bacteriocin pairs under these conditions. We therefore sought to experimentally assess peptide association using split-luciferase complementation assays.

Initial attempts to express bacteriocin-luciferase fusions in *Escherichia coli* under IPTG induction proved unstable, as transformants frequently acquired loss-of-function mutations within the fusion constructs (**Supplementary Fig. 5**). We therefore adapted the assay to a cell-free expression platform, enabling independent synthesis of LgBit-and SmBit-tagged bacteriocins followed by mixing and quantification of NanoLuc reconstitution as a readout of peptide association (**Fig. 4a**).

**Figure 4.**
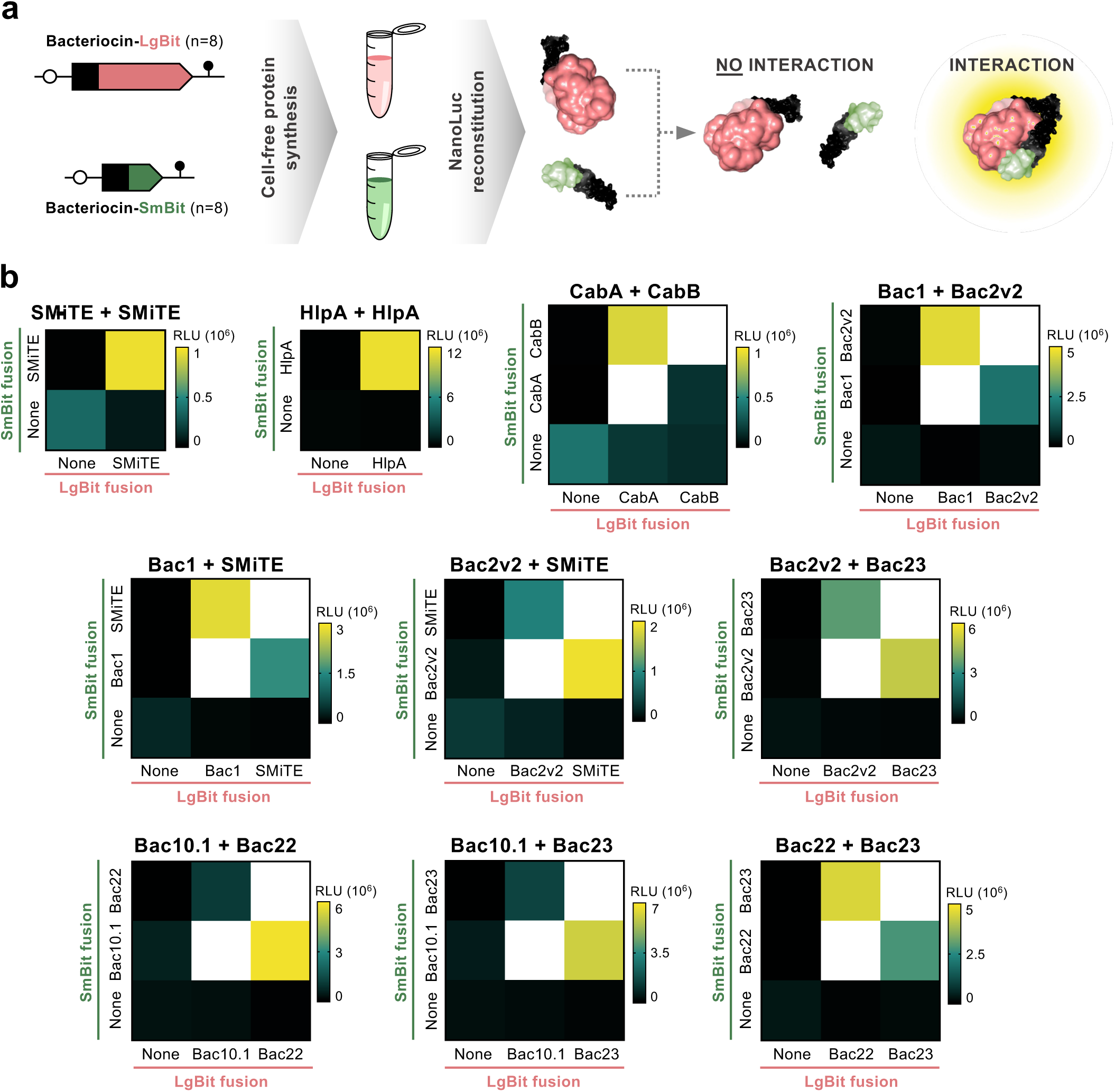
Split-luciferase complementation reveals association between synergistic bacteriocin partners. (a) Representation of the split-luciferase complementation assay. Bacteriocins were synthesized by cell-free protein synthesis as N-terminal fusions to the large (LgBit) or small (SmBit) fragments of NanoLuc luciferase. Fusions were mixed 1:1 after synthesis, and peptide association was quantified by reconstitution of NanoLuc activity and measurement of luminescence. (b) Positive controls for the split luciferase assay included SMiTE, pneumococcal HlpA, and the two-peptide bacteriocin CabA-CabB. The assay was then applied to active bacteriocin pairs identified in *S. mitis* C22, including both intra-locus and inter-locus combinations. Luminescence values were calculated as the area under the curve during the first 5 min following addition of NanoLuc substrate. For clarity, a different scale was used for each pair. Data represents the mean of three biological replicates, each with two technical replicates.

We validated the assay using proteins with known interaction partners. Bacteriocin SMiTE, previously reported to form oligomeric assemblies^16^, produced significantly greater luminescence than all negative controls (**Fig. 4b and Supplementary Table 6**). Likewise, the pneumococcal protein HlpA, which forms homodimers^32^, generated a robust complementation signal consistent with a previous NanoLuc-based interaction study^33^. Finally, the two-peptide bacteriocin CabA-CabB^6^ also produced strong luminescence, confirming that the assay detects association between compatible bacteriocin partners.

We next examined synergistic bacteriocin pairs identified in *S. mitis* C22. Every pair tested generated a detectable complementation signal (**Fig. 4b and Supplementary Table 6**). These included intra-locus associations between Bac1-Bac2v2 (*blp1*) and Bac22-Bac23 (*cab4*), as well as multiple inter-locus associations spanning Bac1-SMiTE (*blp1*-*blp4*), Bac2v2-Bac23 (*blp1*-*cab4*), and Bac10.1-Bac22 and Bac10.1-Bac23 (*blp6*-*cab4*).

To further assess whether peptide association depends on bacteriocin sequence, we selected two robust inter-locus pairs, Bac2v2-Bac23 and Bac22-Bac23, and compared their interactions with scrambled variants. Scrambling either peptide generally abolished luciferase complementation, whereas scrambling both peptides consistently eliminated detectable association, indicating that the interaction depends strongly on the specific peptide sequences (**Supplementary Fig. 6 and Supplementary Table 6**). This effect was observed with both orientations of the NanoLuc fragments, although one exception was observed for Bac2v2_Sc_-SmBit (scrambled) with Bac23-SmBit, which retained a complementation signal that was not significantly different from that of the native pair. Notably, the corresponding reciprocal fusion, Bac2v2_Sc_-LgBit with Bac23-SmBit, did not retain this interaction. These results indicate that peptide association is predominantly sequence-dependent, while also suggesting that physicochemical properties may contribute to association in some contexts.

Together, these findings demonstrate that active bacteriocin pairs consistently exhibit peptide association, supporting a model in which functional two-peptide bacteriocins assemble through selective pairing between compatible peptides.

### Inter-locus bacteriocin pairs enhance competitiveness in co-culture

To determine whether functional bacteriocin pairs identified by cell-free protein synthesis could also function in a native bacterial context, we constructed a bacteriocin-free *S. mitis* strain in which individual bacteriocins could be expressed independently. For that, we selected *S. mitis* G22 because of its high genetic tractability and well-defined bacteriocin repertoire^6,13^. The wild-type G22 genome contains five bacteriocin loci, which we sequentially deleted to generate a bacteriocin-free strain, G22_Naïve_ (**Supplementary Fig. 7a**). We first verified that G22_Naïve_ could be used as a neutral background for bacteriocin expression. Cell-free supernatants (CFS) from wild-type G22 produced a zone of inhibition against *S. pneumoniae*, whereas CFS from G22_Naïve_ showed no detectable inhibitory activity (**Supplementary Fig. 7b**). In contrast, CFS from a G22_Naïve_ derivative expressing SMiTE under the control of the native *blp1* locus promoter displayed potent antimicrobial activity (**Supplementary Fig. 7b**). This result is consistent with our cell-free protein synthesis experiments showing that SMiTE has intrinsic activity against *S. pneumoniae* and does not require a compatible peptide partner for antimicrobial activity.

We next asked whether bacteriocin pairs identified in the cell-free protein synthesis screen could enhance competition when their individual components were produced by distinct bacterial cells. We generated G22_Naïve_ derivatives where individual bacteriocin peptides were integrated in the native *blp1* locus. These strains remained viable and showed no detectable growth defect when bacteriocin expression was induced with exogenous BlpC_1_ (**Supplementary Fig. 8**). We then co-inoculated pairs of G22_Naïve_ derivatives with *S. pneumoniae* D39 and quantified competitive fitness after 12 h as the ratio of *S. mitis* to *S. pneumoniae* recovered from mixed cultures relative to the corresponding inoculum ratio. Production of bacteriocin components that formed active pairs in the cell-free protein synthesis assays increased the competitive index of *S. mitis* for five combinations: Bac1-Bac2, Bac2-Bac23, Bac10.1-Bac22, Bac10.1-Bac23, and Bac22-Bac23 (**Figure 5 and Supplementary Table 7**). Two combinations, Bac1-SMiTE and Bac2-SMiTE, did not produce an increase in competitive index when the two components were produced. However, SMiTE has potent intrinsic antimicrobial activity in the absence of a partner (**Fig. 2a**), which could limit the ability of this assay to resolve additional contributions from peptide pairing.

**Figure 5.**
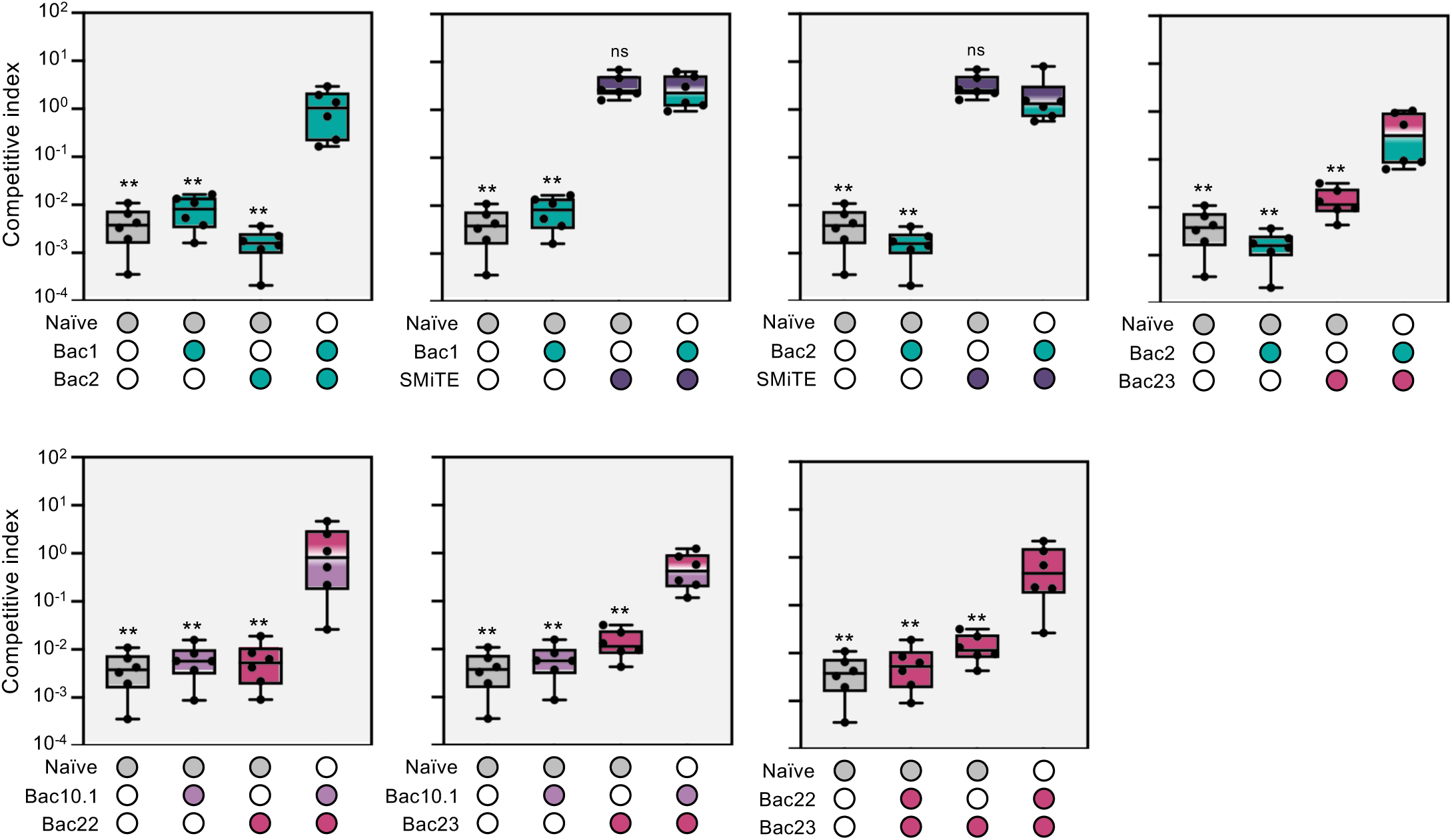
Intra- and inter-locus bacteriocin combinations increase competitiveness in co-culture. *S. pneumoniae* D39 and *S. mitis* (G22_Naïve_ or derivatives) were co-inoculated into fresh C+Y_YB_ supplemented with 1 μM of BlpC_1_ at ∼10^5^ cfu/mL each. When two *S. mitis* strains were included, each was inoculated at ∼5×10^4^ cfu/mL each. *S. pneumoniae* and *S. mitis* cfu/mL were enumerated at 0 h (inoculum) and after 12 h by plating on BA containing tetracyclin or kanamycin, respectively. Competitive indices were calculated as the ratio of *S. mitis* to *S. pneumoniae* recovered after 12 h divided by the corresponding ratio in the initial inoculum. Data represents the mean of three biological replicates, each with two technical replicates. Two-tailed Mann-Whitney *U* test was used for statistical comparison. ** < 0.01, ns not significant. Statistical results are represented in comparison to the condition where two G22_Naïve_ derivatives expressing two different bacteriocins were co-inoculated.

Together, these results demonstrate that bacteriocin components identified as compatible in cell-free protein synthesis assays also functionally cooperate when produced by separate bacterial cells, increasing the competitive fitness of *S. mitis* in co-culture with *S. pneumoniae*. Thus, functional bacteriocin pairing is not restricted to peptides produced within the same cell or encoded at the same genomic locus, but can extend across bacterial strains, providing a mechanism through which bacteriocin interactions may contribute to competition within microbial communities.

## Discussion

Two-peptide bacteriocins are viewed as genetically self-contained systems in which two peptides are encoded within the same operon and co-expressed to generate a compatible antimicrobial pair^18^. Our findings expand this model by showing that functional pairing is not restricted by genomic proximity. By combining comparative genomics, systematic functional screening, transcriptional reporters, peptide-interaction assays, and competition experiments, we demonstrate that bacteriocin components encoded in distant chromosomal loci and even by different strains can form active pairs and contribute to bacterial competition.

Our comparative genomic analysis indicated that streptococcal genomes contain substantial potential for inter-locus bacteriocin pairing. Nearly half of the analysed genomes encoded bacteriocins with (G/A/S)xxx(G/A/S) motifs at multiple chromosomal loci, and positive co-occurrence between bacteriocins encoded at distinct loci were substantially more frequent than negative associations. Although co-occurrence does not establish functional interaction, these patterns indicate that dispersed bacteriocin repertoires are common and that distinct bacteriocin components are frequently mantained within genomes. Systematic functional screening provided direct evidence that this genomic organization can translate into functional diversity, as active combinations occurred more frequently between peptides encoded at different loci than between peptides originating from the same locus. Previous studies have described individual examples of non-cognate pairing between two-peptide bacteriocins^20–22^, and a recently described two-peptide bacteriocin with components encoded approximately 70 kb apart was considered an unusual deviation from the classical operon-based model^34^. Our results suggest that these are not exceptions, but may instead represent a broader property of bacteriocin repertoires. Importantly, not all bacteriocins in our screen were promiscuous. Some peptides displayed restricted partner specificity, whereas others interacted productively with many peptides from different loci and, in some cases, different strains. Thus, bacteriocin repertoires can contain both selective and promiscuous bacteriocin components, allowing for a finite number of bacteriocin genes to generate a substantially larger set of antimicrobial combinations.

A prerequisite for inter-locus pairing to occur in vivo, is for compatible peptides to be available extracellularly at the same time. Our results provide evidence for temporal coordination of genetically dispersed bacteriocins. In one strain, four bacteriocin loci distributed across the chromosome displayed very similar expression dynamics, with transcription peaking during exponential growth, despite being controlled by partially independent signalling pathways. Cross-regulation among the *blp* loci likely contributed to this coordinated expression, whereas *cab* remained specifically responsive to competence signalling under the conditions examined. We cannot exclude additional regulatory links under alternative environmental conditions, particularly given the extensive interplay between competence and bacteriocin regulatory networks described across several streptococcal species. For example, in *S. pneumoniae, blp* and streptococcin E bacteriocins are induced by ComE^5,31,35^, the transcriptional activator that drives expression of early-competence genes; and the two-peptide bacteriocin CibAB is induced by ComX, the alternative sigma factor that drives expression of late-competence genes^36–38^. Additional links have been found in *S. mutans*^39–42^*, S. mitis*^6,13,30^, *S. salivarius*^43^ and *S. sanguinis*^44^, among others, although their regulatory mechanisms are not always completely understood. Importantly, the coordination observed in *S. mitis* C22 should not be interpreted as evidence that all bacteriocin loci are always expressed simultaneously. Rather, bacteriocin production and competence are environmentally regulated processes whose induction can shift with population density, pH, nutrient availability and neighbouring microbial communities^6,28,29,45–48^. Therefore, it is likely that environmental conditions dynamically shape which bacteriocins are expressed and therefore which peptide combinations can form at a given time.

Crucially, the functional consequences of distributed bacteriocin pairing were not restricted to cell-free protein synthesis assays. Deletion of all five bacteriocin loci from *S. mitis* G22, generated a bacteriocin-free strain, from which bacteriocin components could be expressed individually. This provided a defined system to test whether peptide combinations identified in the cell-free protein synthesis screen could enhance bacterial competition when produced in a native host. Co-culture experiments showed that several active combinations increased the competitive fitness of *S. mitis* when the two peptide components were produced by distinct bacteria. These findings demonstrate that functional bacteriocin pairing can occur in a native bacterial context and that compatible peptide components do not necessarily need to be encoded or produced within the same strain to contribute to microbial competition.

The ability of distinct bacterial cells to supply compatible peptide components has broad implications for bacteriocin-mediated competition. Our experiments demonstrate this principle directly for selected combinations in an engineered *S. mitis* background, but they do not establish that such cross-cell pairing occurs naturally between unrelated strains in microbial communities. Nevertheless, this finding provides a basis for considering bacteriocin activity at the community level rather than exclusively as a property of individual genomes or cells. In natural communities, bacteriocin-producing strains can coexist and can respond to pheromones produced by neighbouring cells^6,49^, creating opportunities for temporal overlap in bacterial production. Under such conditions, peptides released by different strains could potentially encounter compatible peptide partners in the extracellular environment. Our previous observation that a mixture of the seven streptococcal strains used here exhibited greater antagonistic activity against *S. pneumoniae* than the individual strains^13^ is consistent with the hypothesis that bacteriocin repertoires are a community-level feature. Still, direct demonstration of bacteriocin pairing across naturally occuring isolates will be required to determine how frequently this occurs in polymicrobial communities.

Our findings also raise the possibility that productive peptide partners may not be restricted to bacteriocins. Host mucosal surfaces contain abundant antimicrobial peptides that coexist with bacteriocins during colonization. If some bacteriocins are capable of forming partnerships with diverse peptides, interaction between bacteriocins and host-derived antimicrobial peptides may also occur. Although speculative, this hypothesis is consistent with previous observations linking host antimicrobial peptides with competence and bacteriocin regulation^50–52^. For example, exposure of *S. pneumoniae* to the human antimicrobial peptide LL-37 induces both competence^53^ and bacteriocin-associated regulatory pathways^54^, suggesting previously unappreciated connections between bacterial competition and host immunity.

The promiscuous pairing of bacteriocin peptides also raises questions about how producer cells avoid self-toxicity. In the classical model, an immunity protein is typically encoded and expressed alongside its cognate bacteriocin, providing immediate protection. Combinatorial pairing introduces a different set of possibilities: a single immunity determinant could protect against multiple complexes^55,56^, whereas alternatively, different peptide combinations could require distinct or partially overlapping immunity mechanisms. Yet, our experiments suggest that dedicated cognate immunity may not be strictly required. Individual bacteriocin peptides were introduced into the G22_Naïve_ background without co-introduction of cognate immunity determinants, and the resulting strain remained viable and showed no growth defect following bacteriocin induction with BlpC. Notably, this was also true for SMiTE, which displayed strong antimicrobial activity by itself against *S. pneumoniae*, indicating that cognate immunity was not required for resistance under these conditions. Several explanations could account for this observation. For example, bacteriocin activity may depend on the presence of a specific receptor in the cell membrane ^57–60^, which could render the producer intrinsically resistant to its own bacteriocins if it lacks such a receptor. Alternatively, endogenous cellular mechanisms or immunity determinants encoded elsewhere in the genome may provide protection. Consistent with this, the seven strains analysed here encode more putative immunity determinants than bacteriocins^13^, including immunity determinants that are not immediately adjacent to any predicted bacteriocin, suggesting that immunity does not always follow a one-to-one organization. Some bacteriocins may also be involved in mechanisms of self-protection^61^. Defining how these diverse mechanisms accommodate combinatorial bacteriocin pairing will be important for understanding the evolutionary stability and ecological consequences of bacteriocin repertoires.

Our findings further suggest a potential mechanism for maintaining bacteriocin diversity during genome evolution. In a strict cognate two-peptide system, changes to one peptide can easily compromise compatibility with its partner, potentially constraining sequence diversification. In contrast, a distributed interaction network provides alternative routes to functionality. Newly acquired or mutated peptides may retain activity if they interact with one more compatible partners already present elsewhere in the genome. Whether this architecture actually facilitates the evolutionary acquisition or retention of novel bacteriocin genes remains to be tested.

Together, these findings establish bacteriocin pairing as a modular process in which antimicrobial potential of a genome is not determined solely by the number of bacteriocin genes it encodes, but also by the combinations that those peptides can form.

## Methods

### Co-occurrence analysis

For co-occurrence analysis of two-peptide bacteriocins among *Streptococcus* spp., a dataset of bacteriocins identified from RefSeq using IIBacFinder was used^9^ (**Supplementary Table 1**). Several filtering steps were applied to this collection. Bacteriocins located near contig edges (within 30 kb of contig boundaries) were excluded. The remaining bacteriocins were then screened and filtered for the presence of conserved (G/A/S)xxx(G/A/S) motifs within the predicted mature peptide sequence. Bacteriocins were assigned to the same bacteriocin loci based on a maximum distance of 30 kb.

Co-occurrence analysis was performed using the filtered bacteriocin dataset for all pairwise bacteriocin combinations found in individual strains. For each pair, the expected number of co-occurences under the assumption of independence was calculated from the prevalence of each bacteriocin across the genome collection, and an observed-to-expected co-occurrence ratio was calculated. Co-occurrences were considered significant when the false-discovery rate (FDR) was < 0.05 and when the pair was observed in at least 10 genomes. Positive and negative co-occurrences were considered when the odds ratio (OR) was > 2 or < 0.5, respectively.

### Bacterial strains and growth conditions

Bacterial strains, primers and peptides used in the study can be found in **Supplementary Table 8**. *S. pneumoniae* was grown at 37 °C in Todd-Hewitt broth with 0.5% yeast extract (THY), *S. mitis* was grown in C+Y_YB_ at 37 °C, and *E. coli* was grown in lysogeny broth (LB) at 37°C with shaking. When appropriate, media was supplemented with 8 μg.mL^-1^ chloramphenicol (*S. mitis*), 300 μg.mL^-1^ (*S. mitis*) or 100 μg.mL^-1^ kanamycin (*E. coli*), 200 μg.mL^-1^ (*S. mitis*) or 100 μg.mL^-1^ (*E. coli*) spectinomycin, and 2 μg.mL^-1^ tetracycline (*S. pneumoniae*).

### Transformation

*S. mitis* C22 and G22 were grown in C+Y_YB_ (pH 7.4) at 37 °C until an OD600 of 0.5, and diluted 1:100 in fresh medium. After reaching OD_600_ of 0.04 to 0.1, cultures were treated with 1 μM of CSP_C22_ and 400 ng.mL^-1^ of linear DNA or 1 μg.mL^-1^ of plasmid DNA. After 2-4 h at 37°C, transformants were recovered in tryptic soy agar plates (TSA) supplemented with 5% sheep blood (BA) and appropriate antibiotics. Transformants were further confirmed by PCR.

CaCl_2_-competent *E. coli* cells were transformed by heat-shock at 42 °C for 40 s, immediately chilled on ice, and recovered in SOC (2% tryptone, 0.5% yeast extract, 10 mM NaCl, 2.5 mM KCl, 10 mM MgCl_2_,10 mM MgSO_4_ and 20 mM glucose; pH 7.0) for 1 h before plating in LB agar with appropriate antibiotics.

### Antimicrobial activity of cell-free produced bacteriocins

Synthetic DNA constructs encoding bacteriocins were previously designed^16^ to include a minimal T7 promoter, optimized ribosome-binding site^62^, codon-optimized mature bacteriocin sequence, and T7 terminator. DNA was synthesized by Twist Biosciences, resuspended in nuclease-free water, and stored at −20 °C.

Cell-free protein synthesis was performed using PUREfrex 2.0 (GeneFrontier) according to the manufacturer’s instructions. Reactions containing 10 nM of DNA (or nuclease-free water as negative control) were incubated at 37 °C for 4 h.

For antimicrobial assays, *S. pneumoniae* D39 was grown in THY until an OD_600_ of 0.5 and diluted to 10^4^ cfu.mL^-1^ in fresh medium containing 1600 U.mL^-1^ catalase (Sigma). For single-bacteriocin treatments, 90 μL of bacterial suspension was dispensed into 384-well plates and treated with 10 μL of cell-free reactions. For bacteriocin pairwise treatments, 5 μL of each bacteriocin reaction was added. Growth was monitored every 30 min for 24 h using the Tecan Infinite 200 Pro. A peptide was considered active if it increased the lag phase by at least 2 h compared to each of the bacteriocins added individually.

### Construction of bacteriocin reporter strains

Promoter regions corresponding to the upstream intergenic sequences of bacteriocin operons were amplified from *S. mitis* C22 genomic DNA. Promoter fragments and plasmid pJB11^6^ were assembled using BsaI-HFv2 and T4 DNA ligase (NEB) and transformed into *E. coli*. Plasmids were propagated, linearized with BamHI-HF (NEB), and transformed into *S. mitis* C22.

### AlphaFold structural modeling of bacteriocin pairs

Structural models were generated for all 295 pairwise combinations of bacteriocins using AlphaFold 3^63^ with default settings through the AlphaFold server (https://alphafoldserver.com). Interface predicted template modeling (ipTM) scores were then extracted from the model with highest ranking confidence score.

### Measurement of bacteriocin expression using luminescent reporters

Reporter strains were grown in C+Y_YB_ until an OD_600_ of 0.5, diluted 1:100 in fresh medium with 340 μg.mL^-1^ D-luciferin (Abcam), and 200 μL were added per well of white 96-well plates with clear bottoms (BRANDplates). D-luciferin was resuspended in ddH_2_O at 15 mg.mL^-1^ and kept in single-use aliquots at −80°C covered from light. The OD_595_ and luminescence (200 gain) were measured every 10 min for 24 h in a BioTek Neo2 Multimode Reader (Agilent).

To test for the effect of pH on natural induction, cultures were pre-grown in the lowest tested pH (6.2), diluted 1:100 in the appropriate pH, and the OD_595_ and luminescence were measured as above.

For pheromone induction assays, cultures were treated with 1 μM of BlpC_1_, BlpC_4_, CSP_C22_, or water and the OD_595_ and luminescence were measured as above.

### Construction of plasmid for split-luciferase complementation assay in *E. coli* lysates

The backbone of plasmid pET28a(+)-8xHis-GB1-SMiTE was amplified and digested with BamHI-HF and HindIII-HF (NEB). DNA fragments containing a BamHI cut site, a BsaI cut site, a GS linker fused to LgBit, an RBS, a PaqCI cut site, a GS linker fused to SmBit, and a HindIII cut site were synthesized (Twist Biosciences) and cut with BamHI-HF and HindIII-HF. Four variants were synthesized: (i) one where GS-LgBit and GS-SmBit have start codons and no BsaI or PaqCI cut site; (ii) one where GS-LgBit has an ATG and a bacteriocin can be fused to GS-SmBit by cutting with PaqCI; (iii) one where GS-SmBit has an ATG and a bacteriocin can be fused to GS-LgBit by cutting with BsaI; and (iv) one where a bacteriocin can be fused to GS-LgBit by cutting with BsaI and another bacteriocin can be fused to GS-SmBit by cutting with SmBit. The backbone and synthetic constructs were ligated and transformed into *E. coli* DH10b to generate plasmids pJB12a (ATG-GS-LgBit and ATG-GS-SmBit), pJB12b (ATG-GS-LgBit and PaqCI-GS-SmBit), pJB12c (BsaI-GS-LgBit and ATG-GS-SmBit) and pJB12d (BsaI-GS-LgBit and PaqCI-GS-SmBit).

Mature bacteriocin sequences were amplified from *S. mitis* G22 (positive control for two-peptide bacteriocins CabA and CabB) and *S. mitis* C22 (remaining bacteriocins), and *hlpA* was amplified from *S. pneumoniae* D39 (positive control). DNAs and pJB12b, pJB12c and pJB12d were digested with BsaI-HFv2 and ligated to create N-terminal fusions with GS-LgBit, and PaqCI to create N-terminal fusions with GS-SmBit. Plasmids were first transformed and propagated in *E. coli* DH10b and then transformed in *E. coli* BL21. Plasmids were confirmed by PCR and Sanger Sequencing (STAB VIDA). The final plasmids contained a pair of bacteriocins fused to LgBit and SmBit which are transcribed as an operon under control of a IPTG-inducible promoter.

### DNAs for in vitro split-luciferase complementation assay

DNAs for each bacteriocin were synthesized (**Supplementary Table 9**) (Twist Biosciences). DNAs were optimized by GeneFrontier to containe a minimal T7 promoter (TAATACGACTCACTATA), a stemloop (GGGAGACCACAACGGTTTCCC), an AT-rich region (TCTAGAAATAATTTTGTTTAACTTTAAG), a ribosome-binding site (AAGGAG), a spacer (ATATACCA), each bacteriocin fused to a GS linker and either LgBit or SmBit (codon-optimized for *E. coli*), and two consecutive stop codons (TAATGA). To enable synthesis of constructs with <300 bp after optimization, a random DNA sequence containing no start codons was added to the 3’ termini. DNAs were eluted in nuclease-free water to 8 nM and kept at −20°C.

### Split-luciferase complementation assay

For split-luciferase complementation assay using lysates of induced *E. coli* BL21, we adapted protocols from previous studies^33,64^. Strains were incubated overnight in LB with kanamycin. Cultures were diluted 1:10 in fresh medium with 0.5 mM IPTG and kanamycin, incubated for 7 h, and adjusted to 0.5 mL at an OD_600_ of 1.0. Cells were centrifuged and supernatants were discarded. Pellets were resuspended in 250 μL of 50 mM Tris-HCl, 0.5 mM EDTA, 5.2 mg.mL^-1^ lysozyme (pH 8.0) and incubated for 90 min at 37°C. Cells were lysed by five cicles of freeze-thawing in liquid nitrogen and a water bath at 37°C. Lysates were added to wells of a white 96-well plate (96 μL per well). Light production by reconstituted NanoLuc was measured using the Nano-Glo Live Cell Assay System (Promega). Just before measuring, the Nano-Glo Live Cell Substrate was diluted 1:20 in Nano-Glo Live Cell Reagent, and 4 μL were added to wells. Luminescence was measured using 120 gain and 10 s integration signal, in a BioTek Neo2 Multimode Reader.

For in vitro split-luciferase complementation assay using cell-free protein synthesis, the bacteriocins fused to LgBit and SmBit, and both subunits alone were synthesized using PUREfrex2.0 at 37°C for 4 h using a final concentration of DNA of 2 nM. Thirty microliters of nuclease-free water was added to wells of a white 96-well plate (Nunc). To test bacteriocin pairs, 1 μL of cell-free protein synthesis reactions of each bacteriocin was added to the wells. As controls, 1 μL of bacteriocins were mixed with 1 μL of in vitro synthesized LgBit or SmBit. Synthesized LgBit was also mixed with LgBit to assess background luminescence in the absence of a bacteriocin pair. Just before measuring, the Nano-Glo Live Cell Substrate was diluted 1:20 in Nano-Glo Live Cell Reagent and 8 μL was added to each well. Luminescence was measured for every 30 s for 15 min, using 60 gain and 10 s integration signal, in a BioTek Neo2 Multimode Reader. Luminescence was quantified by the area under the curve during the first five minutes of the assay, a timing where we observed that light production had plateaued for all pairs.

### Construction of the bacteriocin-free *S. mitis* G22_Naïve_

The bacteriocin-free G22_Naïve_ strain was constructed by sequentially deleting the five bacteriocin loci present in *S. mitis* G22. First, the *blp4* locus was replaced by a *lox66-p3-aph(3)-lox71* cassette (*p3* – constitutive promoter; *aph(3)* – kanamycin resistance gene). Transformants were selected in BA with kanamycin. The *kanR* marker was subsequently excised by transformation with the Cre-expressing plasmid pJB01^13^ at 30°C to allow Cre-mediated recombination between the *lox* sites, generating the Δ*blp4*::*lox72* mutant. The pJB01 plasmid was subsequently cured by incubation at 37°C, exploiting its temperature-sensitive origin of replication.

The *scg* locus was deleted using the same *lox66-p3-kanR-lox71* cassette and *kanR* excision protocol, generating the Δ*blp4*::*lox72* Δ*scg*::*lox72* strain. Bacteriocin loci *cab7* and *slk* were replaced sequentially with *p3-cat* (*cat* – chloramphenicol resistance gene) and *p3-kanR* cassettes, respectively, generating the Δ*blp4*::*lox72* Δ*scg*::*lox72* Δ*cab7*::*p3-chlR* Δ*slk*::*p3-aph(3)* strain.

To generate the final bacteriocin-free background, the *blp1* locus was replaced with a *p3-ant9* cassette (*ant9* – spectinomycin resistance gene), generating G22_Naïve_ (Δ*blp4*::*lox72* Δ*scg*::*lox72* Δ*cab7*::*p3-cat* Δ*slk*::*p3-aph(3)* Δ*blp1*::*p3-ant9*).

To generate strains expressing individual bacteriocin peptides, the same *blp1* locus was instead replaced with constructs containing the corresponding bacteriocin sequence followed by *p3-ant9*, generating G22_Naïve_::*bacteriocin* strains (Δ*blp4*::*lox72* Δ*scg*::*lox72* Δ*cab7*::*p3-cat* Δ*slk*::*p3-aph(3)* Δ*blp1*::*bacteriocin*-*p3-ant9*).

### Growth curves of G22_Naïve_ and derivative strains

Strains were grown in C+Y_YB_ (pH 6.6) until an OD_600_ of 0.5, diluted 1:100 in fresh medium and 198 μL of mixtures were added to clear 96-well plates. Then, 2 μL of water or BlpC_1_ to a final concentration of 1 μM (to induce bacteriocin expression from the *blp1* locus) was added to wells. Growth was monitored every 10 min for 24 h in a BioTek Synergy H1 plate reader.

### Cell-free supernatants and well-diffusion assay

*S. mitis* G22, G22_Naïve_ and derivatives were pre-grow in C+Y_YB_ (pH 6.6) until an OD_600_ of 0.5 and diluted 1:100 in fresh C+Y_YB_ containing 1 μM of BlpC_1_. Cultures were then grown at 37°C until early stationary phase and chilled on ice. Cells were removed by centrifugation and filter-sterilization (0.22 μm), generating cell-free supernatants (CFS). CFS were lyophilized in a vacuum evaporator (LabConco) and resuspended to 1:10 of the initial volume in water (10x CFS).

For well-diffusion assay, *S. pneumoniae* D39 was grown in THY to an OD_600_ of 0.5. Then, a mixture containing 750 μL of culture, 7125 U of catalase, 8 mL pre-warmed THY, and 21 mL of molten tryptic soy broth with 1.5% agar was dispensed in a Petri dish. Wells were created in the agar using the top of 1 mL pipette tips, and 200 μL of 10x CFS or 10x C+Y _YB_ was added to wells. Plates were incubated overnight at 37°C with 5% CO_2_, and growth inhibition halos were measured by two researchers. Plates were imaged using iBright FL1500.

### Competition of S. mitis and S. pneumoniae

*S. pneumoniae* D39-gfp Δ*comGC*^6^ (tetracycline resistant), G22_Naïve_ and derivative mutants (kanamycin resistant) were grown in C+Y_YB_ (pH 6.6) until an OD_600_ of 0.5. For competition experiments, *S. pneumoniae* and G22_Naïve_ were inoculated at a 1:1000 dilution (aproximately 10^5^ cfu/mL of each) in fresh C+Y_YB_ containing 1 μM of BlpC_1_. In cases where two strains of *S. mitis* were co-inoculated with *S. pneumoniae*, aproximately 5×10^4^ cfu/mL of each *S. mitis* strain was used. The cell counts of *S. pneumoniae* and *S. mitis* at 0 h (inoculum) and after 12 h of incubation were enumerated by plating in BA with tetracycline or kanamycin, respectively. Competitive indices were calculated as the ratio of *S. mitis* to *S. pneumoniae* recovered after 12 h divided by the corresponding ratio in the initial inoculum.

### Statistics

Statistical analyses were carried out using Python 3.11 or GraphPad Prism 10. For co-occurrence analysis of bacteriocin pairs, the Fisher’s exact test (scipy.stats) was used, and *P* values were adjusted for multiple comparisons using the Benjamini-Hochberg FDR (statsmodels). For split-luciferase assays, one-way ANOVA with Dunnett’s multiple comparisons test was used. For comparison of ipTM scores of AlphaFold models and competitive indices, the two-tailed Mann-Whitney *U* test was used. For comparison of inhibition halo sizes from well-diffusion assays, the two-tailed students *t*-test with Benjamini and Hochberg correction for FDR was used. Results were considered significant when *P* or FDR < 0.05.

## Author contributions

J.L., J.B. and R.S.-L. contributed to the concept and design of the study. R.S.-L. contributed with reagents and materials. Data acquisition was performed by J.L. and J.B.. Data was interpreted by J.L., J.B. and R.S.-L.. The manuscript was written by J.L., J.B. and R.S.-L.. All authors critically revised and approved the final version of the manuscript.

