## Supplementary Figures for "Multi-locus pairing expands the functional repertoire of two-peptide bacteriocins"

### Supplementary Figure 1

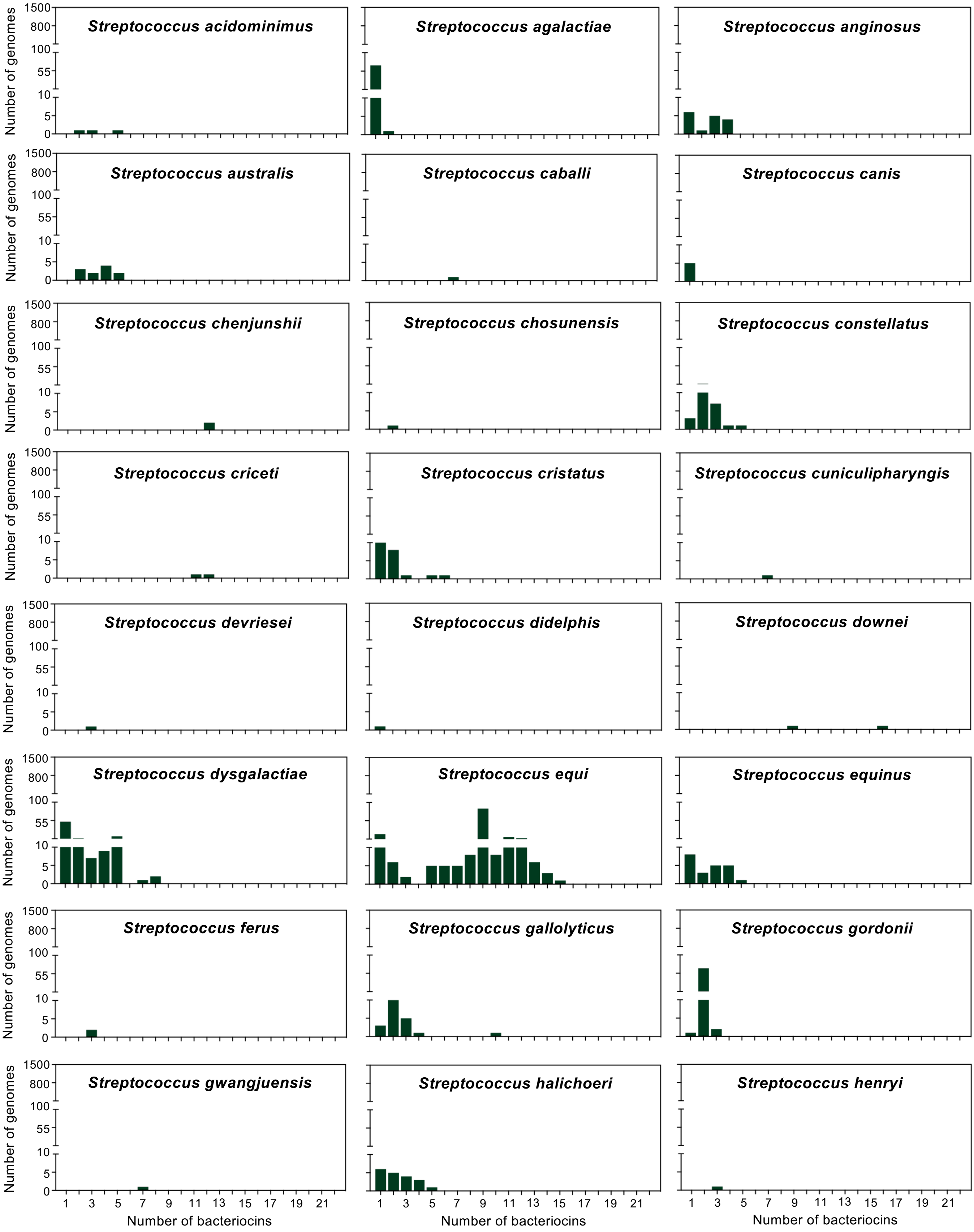


(continues next page)

#### Supplementary Figure 1

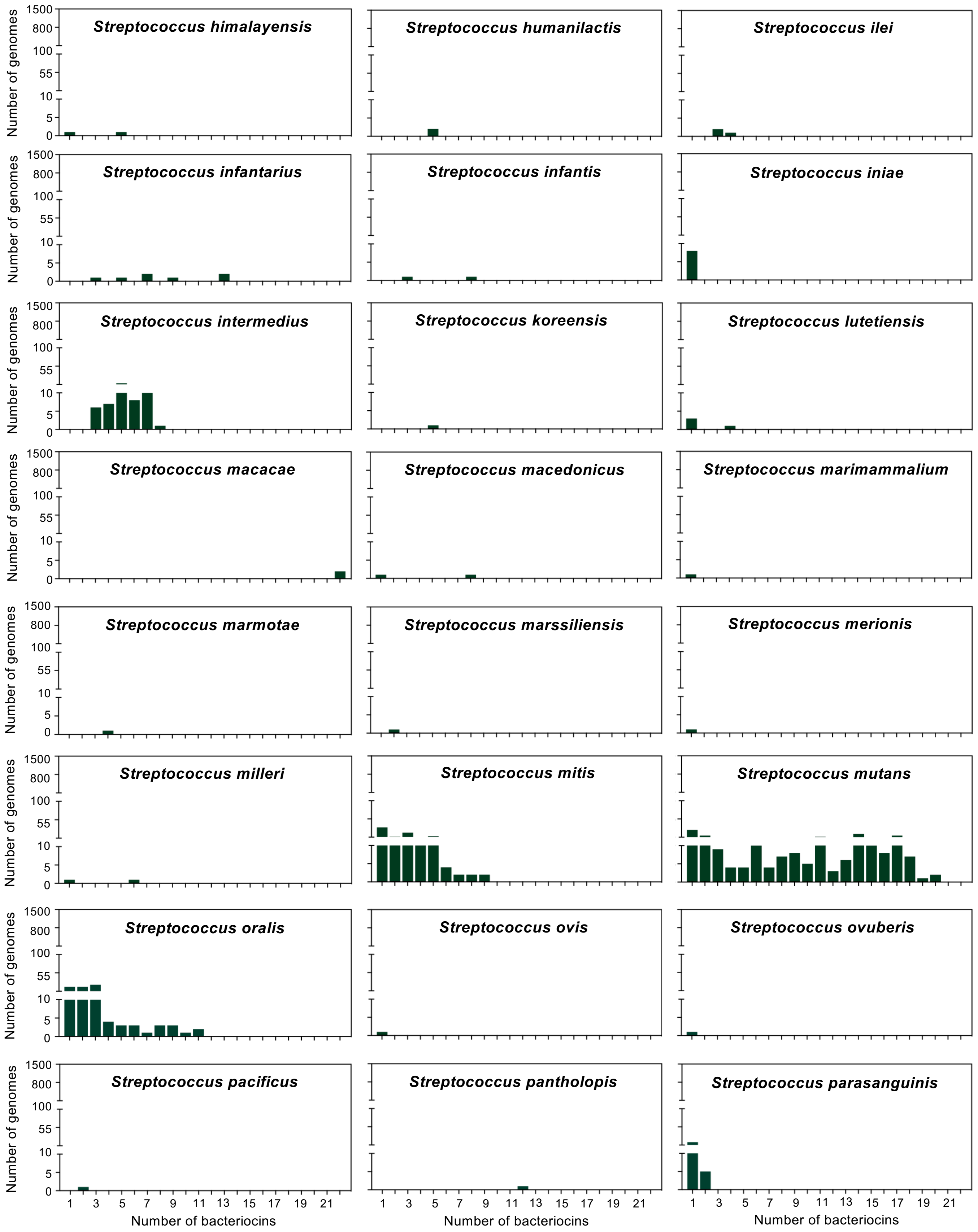


(continues next page)

#### Supplementary Figure 1

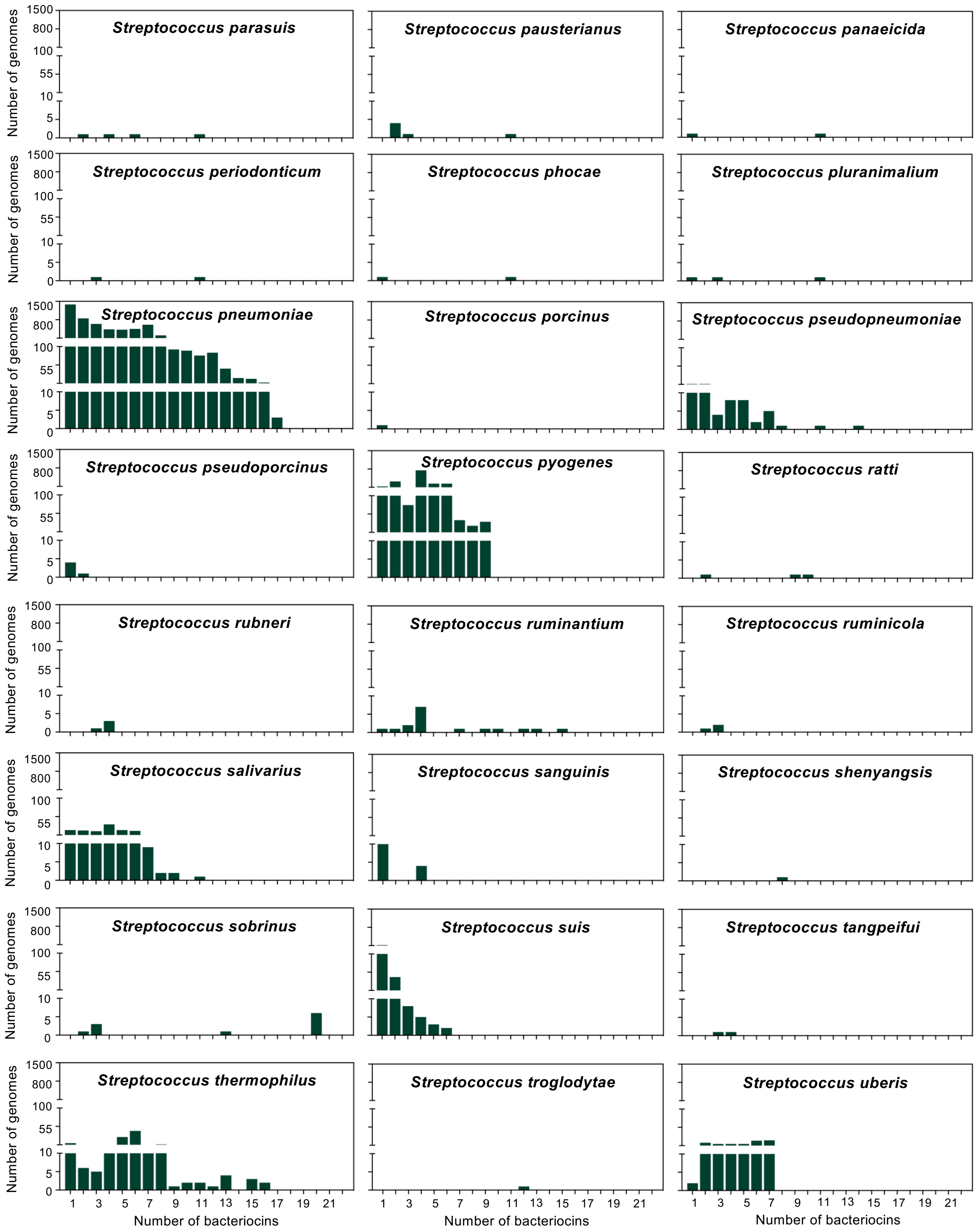


(continues next page)

### Supplementary Figure 1

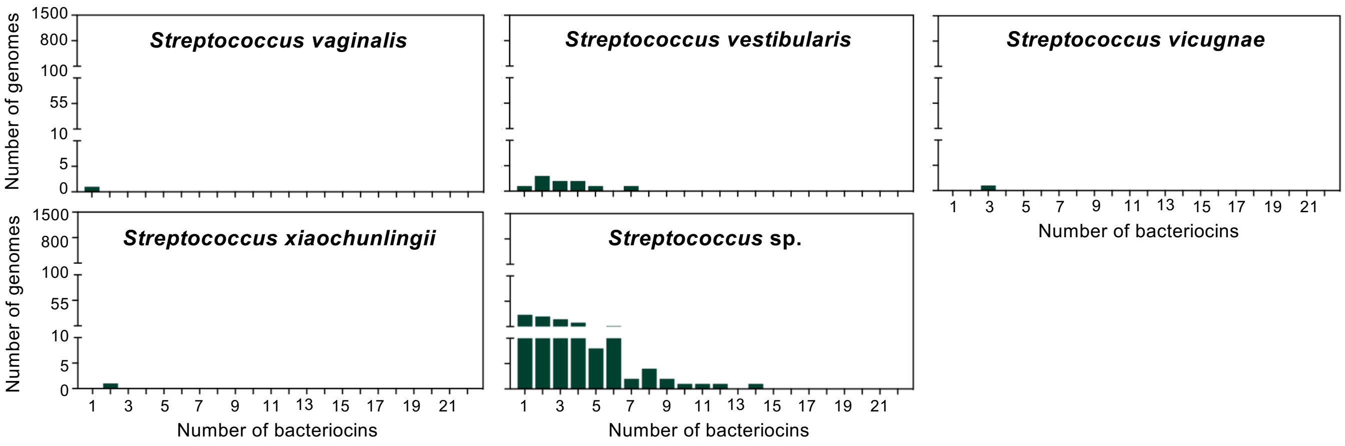


**Supplementary Figure 1. Distribution of the number of bacteriocins in streptococcal genomes.** Data represents results from Figure 1c at a species-specific level. Bacteriocins were filtered to exclude peptides located within 30 kb of contig edges. Bacteriocins containing (G/A/S)xxx(G/A/S) motifs in their predicted mature sequences were retained.

### Supplementary Figure 2

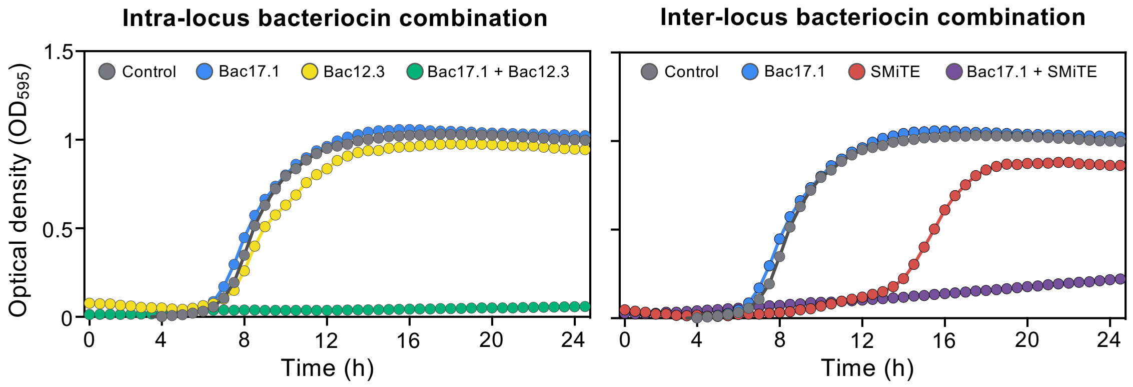


**Supplementary Figure 2. Intra-locus and inter-locus bacteriocin combinations are active against *S. pneumoniae*.** Mature bacteriocins were produced by cell-free protein synthesis and tested individually and in pairwise combinations against *S. pneumoniae* D39. Data represents the mean of 3 biological replicates. Examples of effect of intra-locus (left) and inter-locus (right) bacteriocin combinations from Figure 2a.

### Supplementary Figure 3

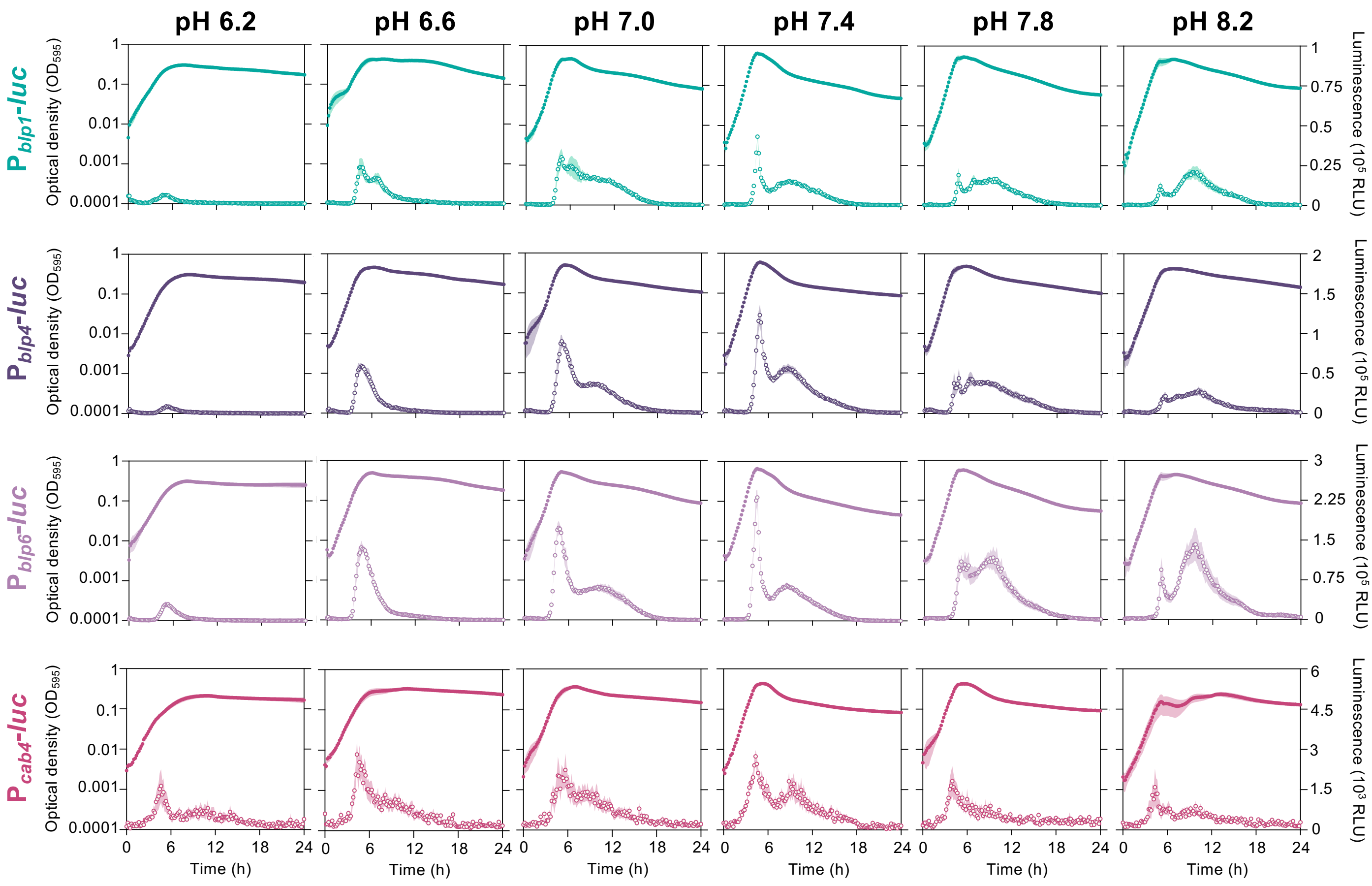


**Supplementary Figure 3. Impact of extracellular pH on the expression of bacteriocin loci of *Streptococcus mitis* C22.** Temporal expression of the four bacteriocin loci (*blp1, blp4, blp6,* and *cab4*) during growth under different media pH (6.2 to 8.2). Promoter activity was monitored using firefly luciferase transcriptional reporters fused to the promoter region of each bacteriocin locus. Filled and open circles represent the optical density and luminescence, respectively. Data represents the mean ± s.d. from three biological replicates, each with three technical replicates.

### Supplementary Figure 4

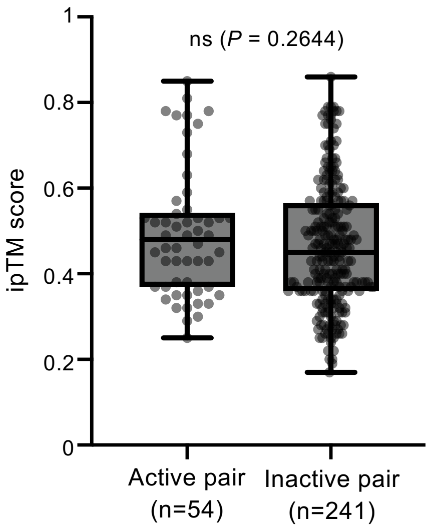


**Supplementary Figure 4. Interface predicted template modeling (ipTM) score of strucutral models of bacteriocin pairs.** Pairwise bacteriocin combinations were modeled using AlphaFold 3 and the ipTM was calculated. Differences between ipTM values of active and inactive bacteriocin pairs were compared using the two-tailed Mann-Whitney *U* test. ns not significant.

### Supplementary Figure 5

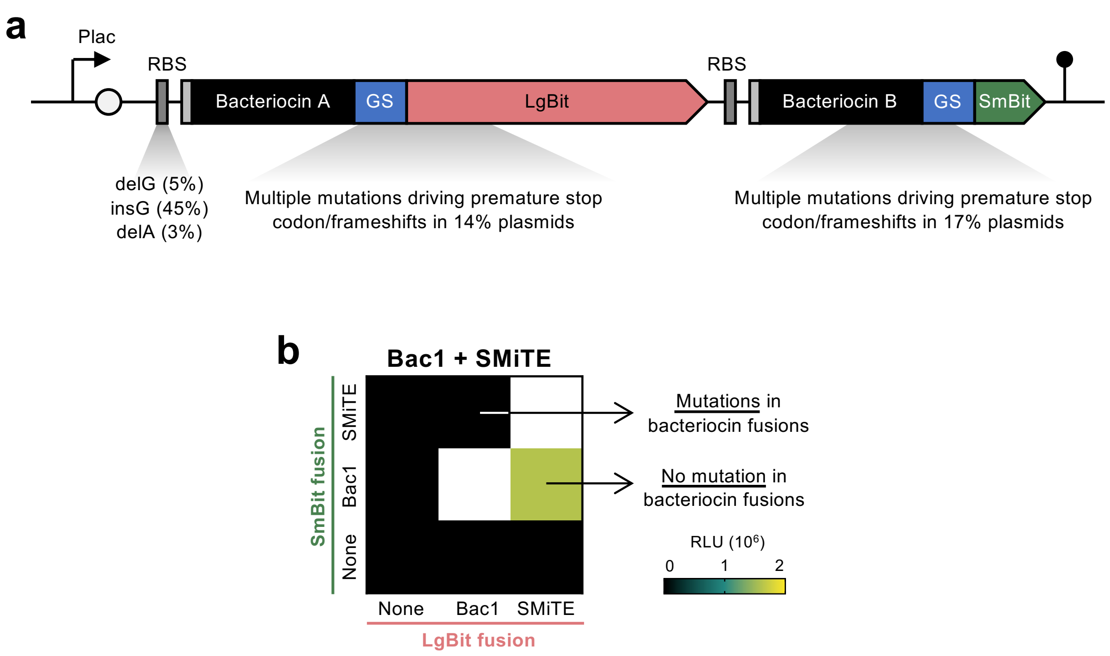


**Supplementary Figure 5. Split-luciferase assays using *Escherichia coli* lysates.** (a) Accumulation of mutations in split-luciferase plasmids. (b) Interaction of Bac1 and SMiTE probed by split-luciferase using *E. coli* lysates after induction with IPTG.

### Supplementary Figure 6

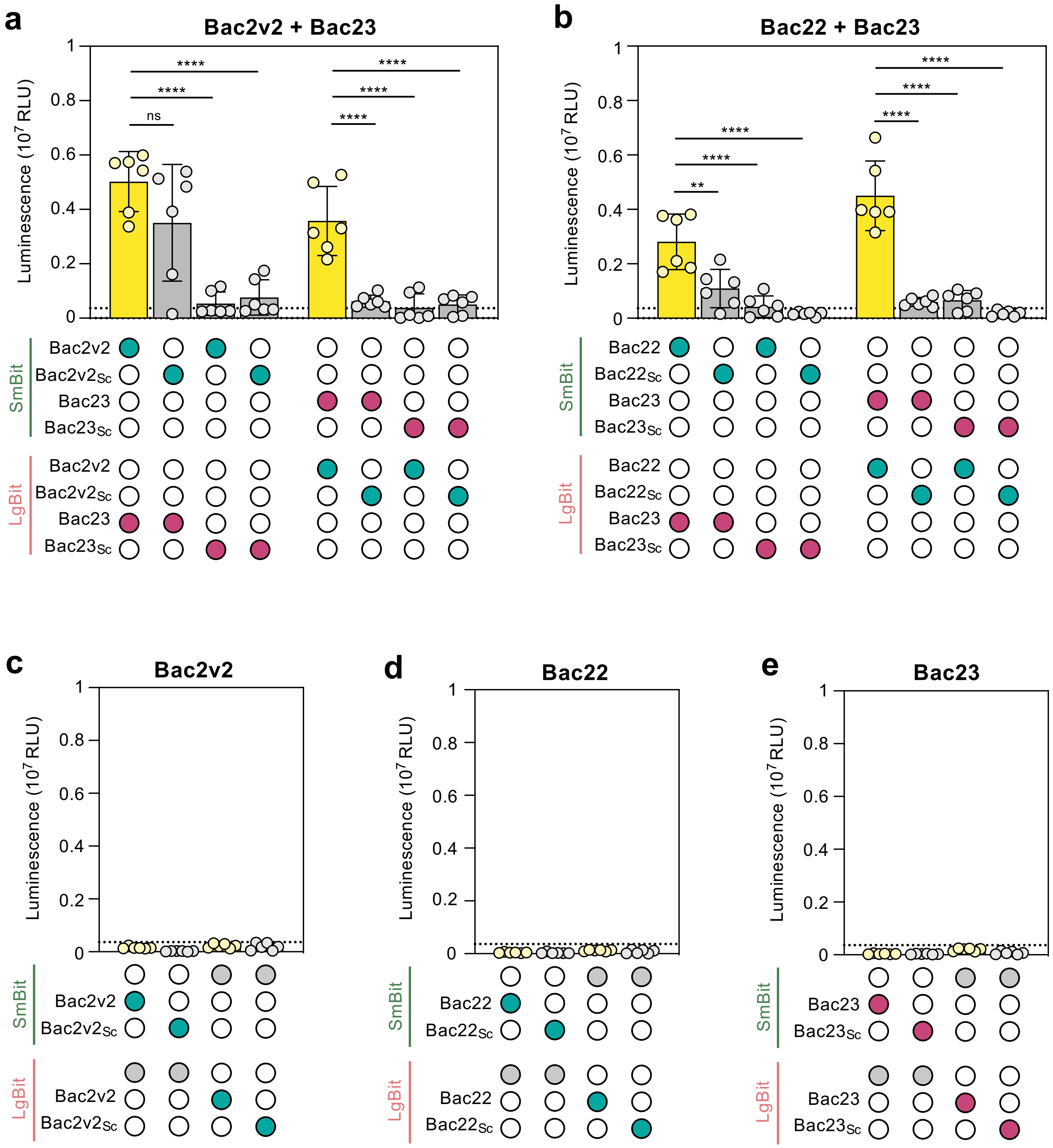


**Supplementary Figure 6. Scrambling bacteriocin sequences decreases split-luciferase complementation.** Wild-type and scrambled (Sc) bacteriocins were synthesized by cell-free protein synthesis as N-terminal fusions to the large (LgBit) or small (SmBit) fragments of NanoLuc luciferase. Fusions were mixed 1:1 after synthesis, and peptide association was quantified by reconstitution of NanoLuc activity and measurement of luminescence. Association of (a) Bac2v2 and Bac23 and of (b) Bac22 and Bac23 fusion variants. (c-e) Controls for Bac2v2, Bac22, and Bac23. Dotted line represents the mean luminescence of the negative control (LgBit mixed with SmBit). Data represents the mean ± s.d. of three biological replicates, each with two technical replicates. One-way ANOVA with Dunnett's multiple comparisons test was used for statistical comparison. ** < 0.01, **** < 0.0001, ns not significant.

#### Supplementary Figure 7

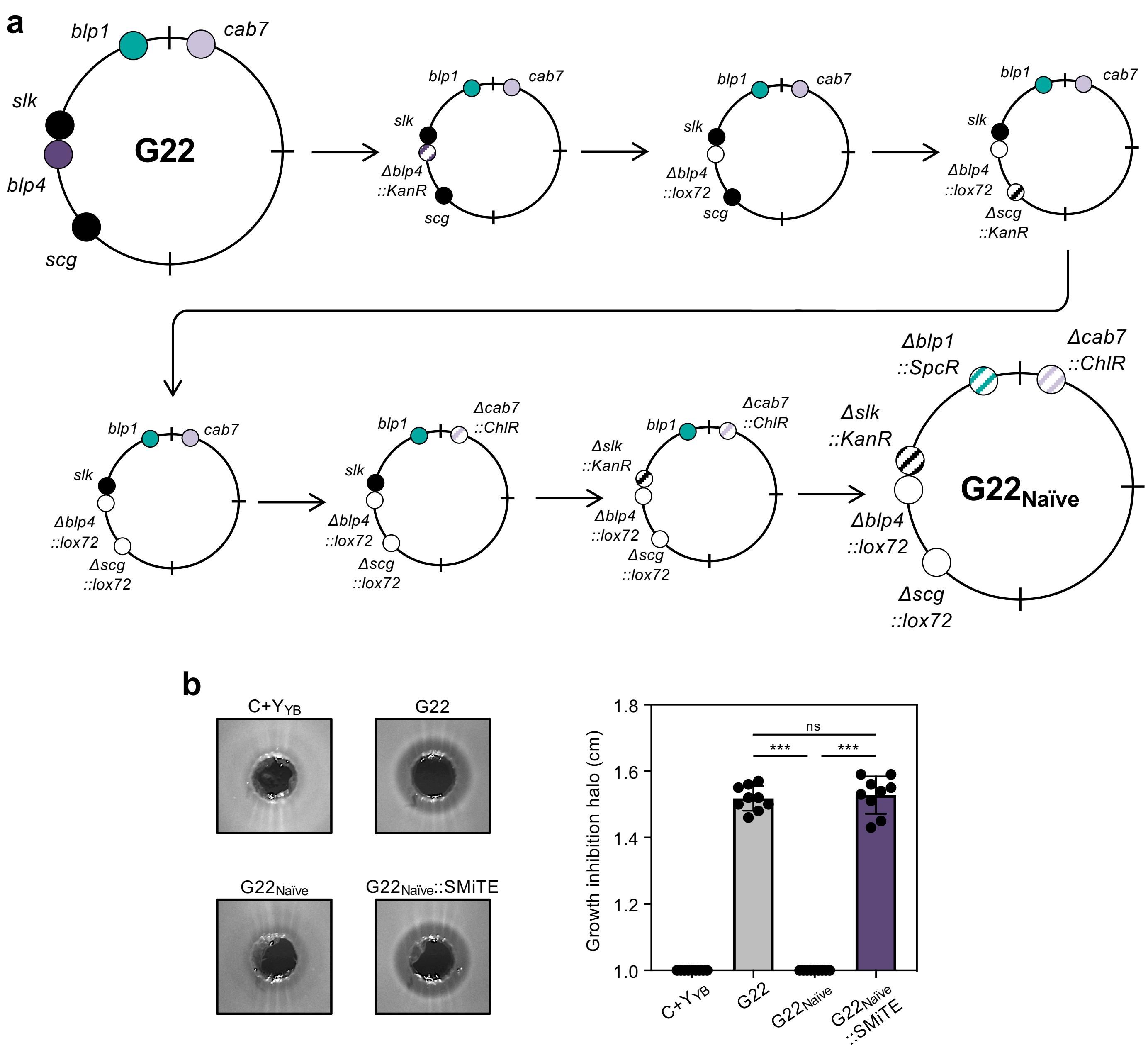


**Supplementary Figure 7. Construction of the bacteriocin-free *S. mitis* G22$_{Naïve}$ and its derivatives.** (a) Sequential genetic manipulations for the construction of G22$_{Naïve}$. (b) Representative images (left panel) and quantification (right panel) of the inhibitory activity of ten-fold concentrated cell-free supernatants of C+Y$_{YB}$, G22, G22$_{Naïve}$ and G22$_{Naïve}$::SMiTE. Each halo was measured by two independent researchers. Each data point represents the mean of each researcher's measurement. Data represents the mean ± s.d. of three biological replicates, each with three technical replicates. Two-tailed students *t*-test with Banjamini and Hochberg correction for FDR was used for statistical comparison. *** < 0.001, ns not significant.

### Supplementary Figure 8

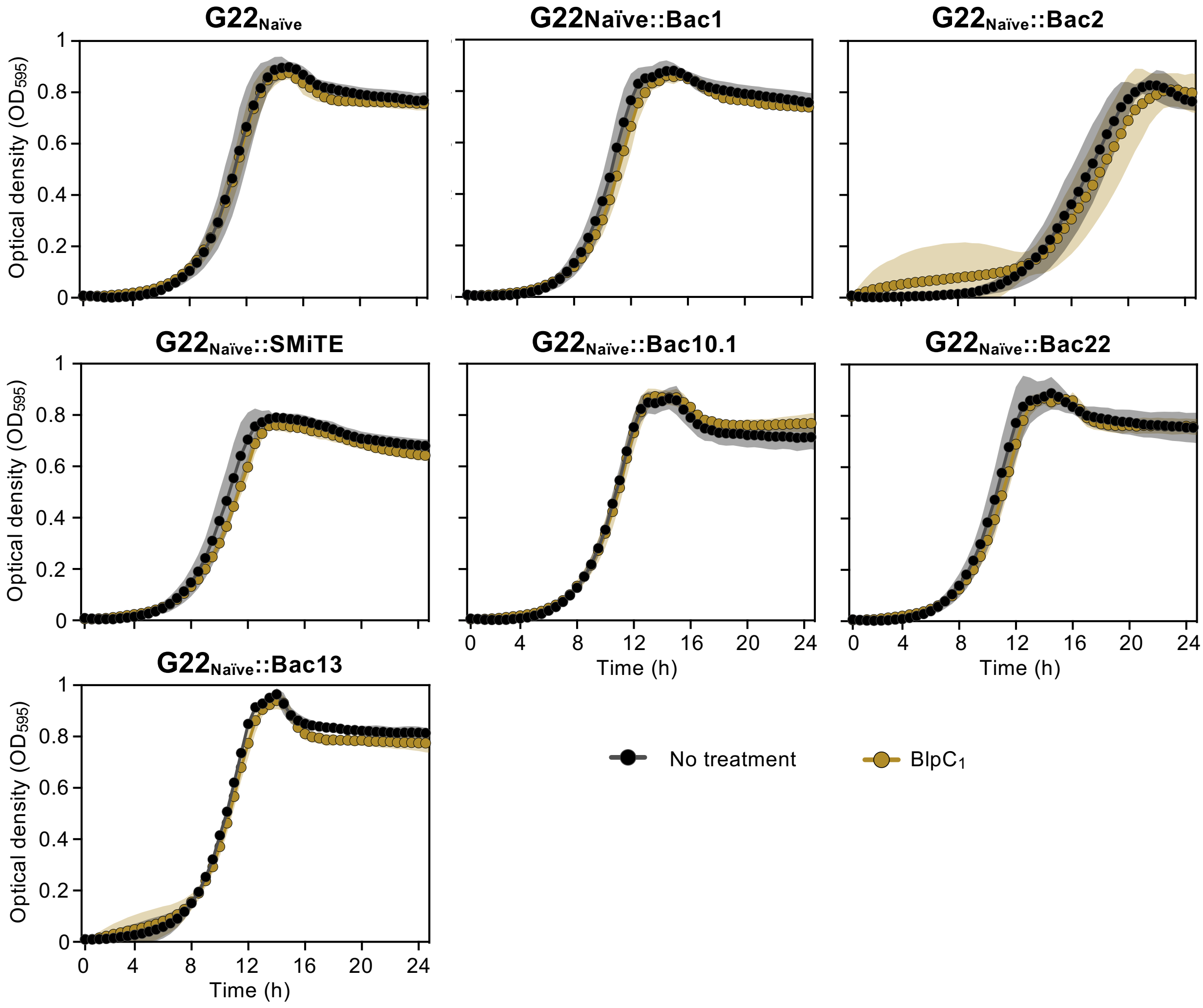


**Supplementary Figure 8. Growth curves of the bacteriocin-free *S. mitis* G22$_{\text{Naïve}}$ and its derivatives.** Strains were grown in C+Y$_{YB}$ pH 6.6 until an $OD_{600}$ of 0.5, diluted 1:100 in fresh medium and 198 µL of mixtures were added to clear 96-well plates. Then, 2 µL of water (no treatment) or $BlpC_1$ to a final concentration of 1 µM (to induce bacteriocin expression from the *blp1* locus) was added to wells. Growth was monitored every 10 min for 24 h in a BioTek Synergy H1 plate reader. Data represents the mean ± s.d. of three biological replicates, each with two technical replicates.
